# Structure-based Classification of Tauopathies

**DOI:** 10.1101/2021.05.28.446130

**Authors:** Yang Shi, Wenjuan Zhang, Yang Yang, Alexey Murzin, Benjamin Falcon, Abhay Kotecha, Mike van Beers, Airi Tarutani, Fuyuki Kametani, Holly J. Garringer, Ruben Vidal, Grace I. Hallinan, Tammaryn Lashley, Yuko Saito, Shigeo Murayama, Mari Yoshida, Hidetomo Tanaka, Akiyoshi Kakita, Takeshi Ikeuchi, Andrew C. Robinson, David M.A. Mann, Gabor G. Kovacs, Tamas Revesz, Bernardino Ghetti, Masato Hasegawa, Michel Goedert, Sjors H.W. Scheres

**Affiliations:** MRC Laboratory of Molecular Biology, Cambridge, UK; Thermo Fisher Scientific, Eindhoven, The Netherlands; Department of Brain and Neurosciences, Tokyo Metropolitan Institute of Medical Science, Tokyo, Japan; Department of Pathology and Laboratory Medicine, Indiana University School of Medicine, Indianapolis, IN, USA; Department of Neurodegenerative Disease and Queen Square Brain Bank for Neurological Disorders, UCL Queen Square Institute of Neurology, London, UK; Department of Neuropathology, Tokyo Metropolitan Geriatric Hospital and Institute of Gerontology, Tokyo, Japan; Molecular Research Center for Children’s Mental Development, United Graduate School of Child Development, University of Osaka, Osaka, Japan; Institute for Medical Science of Aging, Aichi Medical University, Nagakute, Japan; Department of Pathology, Brain Research Institute, Niigata University, Asahimachi, Chuo-ku, Niigata, Japan; Department of Molecular Genetics, Brain Research Institute, Niigata University, Asahimachi, Chuo-ku, Niigata, Japan; Clinical Sciences Building, University of Manchester, Salford Royal Foundation Trust, Salford, UK; Tanz Centre for Research in Neurodegenerative Diseases and Department of Laboratory Medicine and Pathobiology, University of Toronto, Toronto, Canada; Institute of Neurology, Medical University of Vienna, Vienna, Austria

## Abstract

Ordered assembly of the tau protein into filaments characterizes multiple neurodegenerative diseases, which are called tauopathies. We previously reported that by electron cryo-microscopy (cryo-EM), tau filament structures from Alzheimer’s disease (1,2), chronic traumatic encephalopathy (CTE) (3), Pick’s disease (4) and corticobasal degeneration (CBD) (5) are distinct. Here we show that the structures of tau filaments from typical and atypical progressive supranuclear palsy (PSP), the most common tauopathy after Alzheimer’s disease, define a previously unknown, three-layered fold. Moreover, the tau filament structures from globular glial tauopathy (GGT, Types I and II) are similar to those from PSP. The tau filament fold of argyrophilic grain disease (AGD) differs from the above and resembles the four-layered CBD fold. The majority of tau filaments from aging-related tau astrogliopathy (ARTAG) also have the AGD fold. Surprisingly, tau protofilament structures from inherited cases with mutations +3/+16 in intron 10 of *MAPT*, the microtubule-associated protein tau gene, are identical to those from AGD, suggesting that a relative overproduction of four-repeat tau can give rise to the AGD fold. Finally, tau filament structures from cases of familial British dementia (FBD) and familial Danish dementia (FDD) are the same as those from Alzheimer’s disease and primary age-related tauopathy (PART). These structures provide the basis for a classification of tauopathies that also allows identification of new entities, as we show here for a case diagnosed as PSP, but with abundant spherical 4R tau inclusions in limbic and other brain areas. The structures of the tau fold of this new disease (Limbic-predominant Neuronal inclusion body 4R Tauopathy, LNT) were intermediate between those of GGT and PSP.

## INTRODUCTION

Six tau isoforms are expressed in the adult human brain; three isoforms have three microtubule-binding repeats (3R) and three isoforms have four repeats (4R) (6). Based on the isoforms that form the abnormal filaments, tauopathies can be divided into three groups. In PART, Alzheimer’s disease, FBD, FDD and CTE, a mixture of 3R+4R tau isoforms is present in the filaments; 3R tau is found in Pick’s disease; whereas 4R tau isoforms are present in the filaments of PSP, GGT, CBD, AGD and ARTAG. The *H1/H1* haplotype of *MAPT* is a genetic risk factor for 4R tauopathies in some populations (7,8). Dominantly inherited mutations in *MAPT* cause frontotemporal dementias, with tau filaments made of either 3R, 4R or 3R+4R isoforms (9).

We previously showed that Alzheimer’s disease, CTE, Pick’s disease and CBD are each characterized by a different tau protofilament fold (1–5), whereas PART filaments are identical to those from Alzheimer’s disease (10). To expand our knowledge of the diversity of tau filaments in human disease, we determined the structures of tau filaments from the brains of individuals with typical and atypical PSP, GGT (Types I and II), AGD, ARTAG and *MAPT* intron 10 mutations (+3 and + 16), as well as from the brains of individuals with FBD and FDD (**Extended Data Figures 1–3, Extended Data Tables 1-4**).

## RESULTS

First, we examined filaments from PSP, the most common tauopathy after Alzheimer’s disease, and belonging to the group of sporadic frontotemporal lobar degeneration disorders (FTLD-tau). Clinically, typical PSP (Richardson’s syndrome or PSP-RS) is characterised by frequent falls, problems with balance, bulbar dysfunction, cognitive impairment and supranuclear gaze palsy (11,12). It is defined by subcortical neurofibrillary tangles and neuropil threads, together with the presence of tufted astrocytes (13,14). In addition, oligodendroglial coiled bodies and diffuse cytoplasmic tau immunoreactivity are present. Atypical forms of PSP are distinguished by total tau load and specific vulnerability patterns of brain regions (14). We determined the cryo-EM structures, with resolutions up to 2.7 Å, of tau filaments from the frontal cortex and thalamus of three cases of PSP-RS, the putamen and frontal cortex of two cases of PSP with a predominant frontal presentation (PSP-F), the frontal cortex of one case of PSP with predominant parkinsonism (PSP-P) and the frontal cortex of a case of PSP with a predominant presentation of corticobasal syndrome (PSP-CBS).

In PSP-RS, PSP-CBS, PSP-P and PSP-F case 1, we observed tau filaments made of a single protofilament with an ordered core that spans residues 272-381, thus comprising the end of repeat 1, all of repeats 2-4, as well as part of the C-terminal domain. Although this is essentially the same region as that seen in the CBD core, it adopts a markedly different conformation, which we named the PSP fold (**Figure 1a**). Tau filament structures were identical between typical and atypical cases of PSP, consistent with the view that the initiating sites of tau pathology are similar in clinical subtypes and that these subtypes are distinguished by propagation patterns. When multiple tau seeds are found in a PSP brain (15), this may be indicative of copathology.

**Figure 1:**
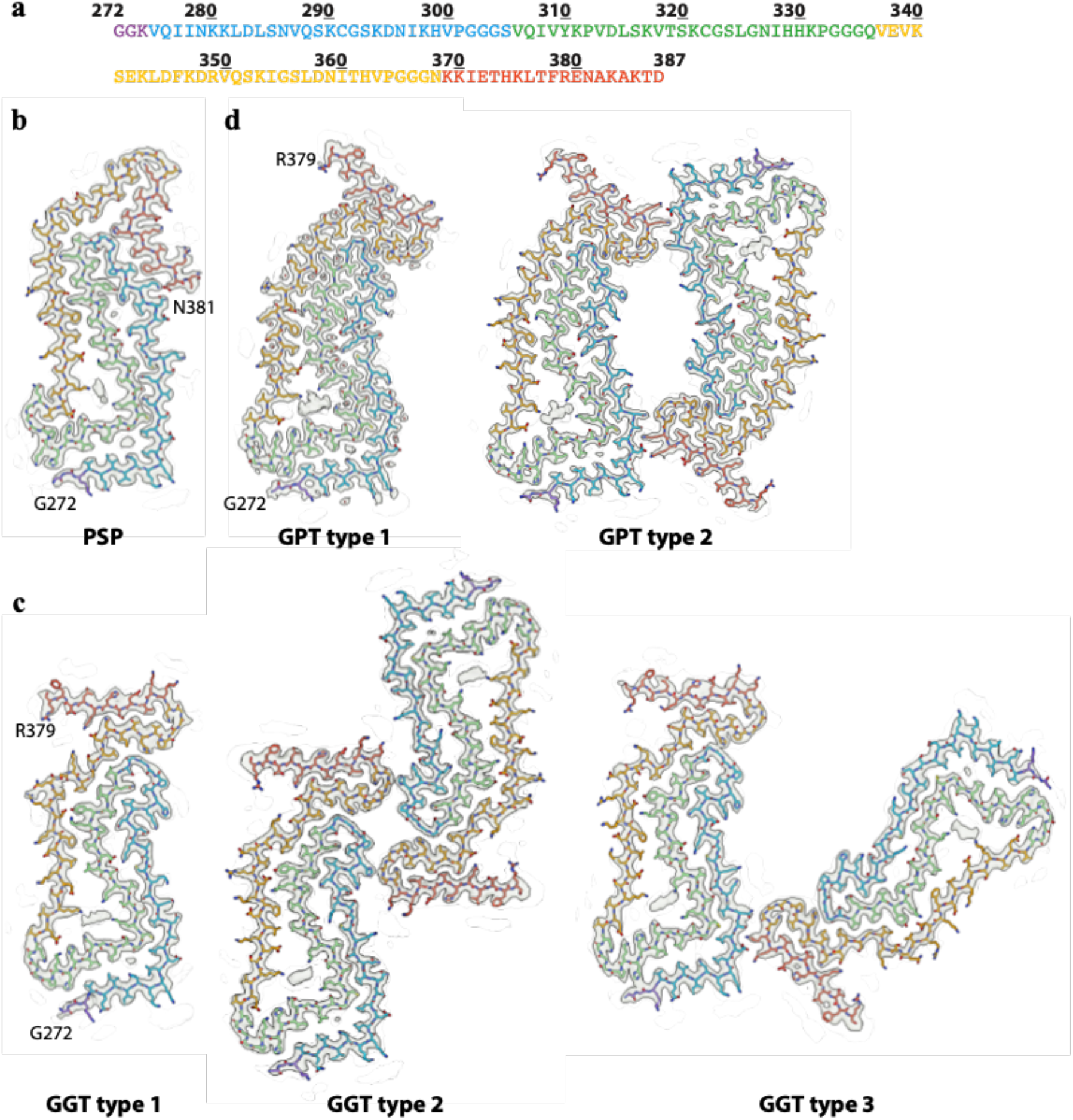
Three-layered 4R tau filament structures. **(a),** Amino acid sequence of the cores of PSP, GPT type 1, GPT type 2, GGT type 1, GGT type 2 and GGT type 3 tau filaments. Residues in R1 are coloured purple; residues in R2 light blue; residues in R3 green; residues in R4 gold; residues in the C-terminal domain dark orange. **(b),** Cryo-EM density map (in transparent grey) and atomic model for the PSP filaments. **(c),** As in (b), for GGT type 1, GGT type 2 and GGT type 3 filaments; **(d)** As in (b), for GPT type 1 and GPT type 2 filaments.

In the PSP fold, repeats 2-4 form a 3-layer meander turning at the conserved PGGG motifs at the end of each tau repeat. R3 forms the central layer, which bends at G323 into a near-right angle. R2 packs against the entirety of R3, following the outside of the bend at G323. The R2-R3 interface contains two small cavities containing additional non-proteinaceous densities. One, between N279 and G323, and surrounded by hydrophobic side chains, is probably caused by molecules of a predominantly non-polar nature, whereas the other, next to the salt bridge between K294 and D314, is most likely a solvent molecule. Most of R4 packs against the other side of R3. At the R3-R4 interface, there is a larger cavity between the positively charged K317, K321 and K340, containing a bigger additional density, presumably corresponding to anionic molecules. This cavity is flanked by the negatively charged E338 and E342, reducing the net positive charge of the cavity to +1 per rung. The rest of the R3-R4 interface is of mixed chemical nature and contains hydrophobic interactions and a salt bridge between K311 and D348. The chain makes another hairpin turn at the PGGG motif of R4 with the extra C-terminal segment forming a short fourth layer covering the end of R2. On the outside of the PSP core, there are a few additional densities that were conserved between cases, the most prominent being next to K280-K281 and H362.

We then examined tau filaments from GGT. Like PSP, GGT is a sporadic 4R tauopathy that belongs to the spectrum of FTLD-tau diseases (16–18). It is divided into Types I-III, with GGT Type I (GGT-I) being characterized by frontotemporal dementia and abundant Gallyas-Braak silver-positive globular oligodendroglial tau inclusions. In GGT Type II (GGT-II), the motor cortex and spinothalamic tract are predominantly affected, whereas GGT Type III (GGT-III) consists of a combination of the clinical features of GGT-I and GGT-II, together with abundant Gallyas-Braak silver-negative globular astroglial inclusions. We determined the cryo-EM structures of tau filaments from the frontal cortex of previously described cases of GGT-I [case 1 in (19)] and GGT-II [case 1 in (20)], with resolutions up to 2.9 Å. Filaments from cases with GGT-I and GGT-II show a common, previously unknown three-layered protofilament fold, spanning residues 272-379 (**Figure 1b**). For GGT-I, we observed three different filament types: type 1 filaments consist of a single protofilament with the GGT fold; type 2 filaments pack two protofilaments with approximate 21 screw symmetry; type 3 filaments pack two protofilaments in an asymmetrical manner. For GGT-II, we only observed filaments of types 2 and 3. We were unable to solve the structures of tau filaments from the frontal cortex of a previously described case with GGT-III [case 1 in (21)], because of an absence of twist in the imaged filaments. This suggests that GGT-III may be a different disease from GGT-I and GGT-II.

The GGT fold comprises essentially the same residues as the PSP fold and has a similar three-layered arrangement of repeats 2-4. Like in the PSP fold, the chain turns at the conserved PGGG motifs, but each turn has a different conformation compared to its PSP counterpart. In addition, the C-terminal domain points in the opposite direction compared to the PSP fold and packs against the end of R4 in a hairpin-like structure that is almost identical to the equivalent part of the Pick fold. The only common substructure of the GGT and PSP folds comprises R2 residues 273-285 and R3 residues 322-330, extending to the additional densities in the small internal cavity between N279 and G323 and outside K280/K281. The cavity at the R3-R4 interface is bigger in the GGT than in the PSP fold and it contains a larger non-proteinaceous density, surrounded by K317, K321 and K340. The cavity’s net positive charge is also greater, as E338 forms a salt bridge with K331.

The protofilament interface in the asymmetric GGT type 3 filaments is formed by residues 283-286 of one protofilament and 367-370 of the other and is stabilised by electrostatic interactions between D283 and K369/K370. In the symmetric GGT type 2 filaments, two small interfaces, between H299 and G366/G367, are stabilised on both sides by additional densities, likely corresponding to non-proteinaceous anionic molecules, between K290 of one protofilament and K369/K370 of the other.

Although the structures of tau filaments from the six cases of typical and atypical PSP described above were identical, we observed a different tau fold for filaments from the frontal cortex of case PSP-F2 (**Figure 1c**). Since this fold resembled both the GGT and PSP folds, we named it the GGT-PSP-Tau, or GPT, fold. We observed two types of GPT filaments: type 1 filaments comprise a single protofilament; type 2 filaments are made of two opposing protofilaments that are related by approximate 21 screw symmetry and create a large, solvent-filled cavity. Histologically, case PSP-F2 resembled cases with abundant spherical, 4R tau-immunoreactive, basophilic neuronal inclusions in limbic and other brain regions (**Extended Data Figure 4**) (22–24).

For PSP-F2, we used a Krios G4 electron microscope with a cold field-emission gun, a Selectris X energy filter and a Falcon-IV camera. The same microscope was used previously for structure determination to atomic resolution (1.22 Å) of an apoferritin test sample (25). Images from this microscope allowed classification into two alternative main chain conformations, one of which led to a reconstruction with a resolution of 1.9 Å **(Extended Data Figure 5)**. This map allowed us to model 19 ordered water molecules per asymmetric unit, but it failed to provide further insights into the nature of the constituent molecules of the additional densities, probably because they did not follow the helical symmetry and were averaged out in the reconstruction process.

The GPT fold comprises the same residues as the PSP and GGT folds and has a similar three-layered structure. The GPT fold has two large substructures of closely similar conformations compared to the GGT fold: the hairpin formed by residues 356-378, as well as residues 273-294 and 312-346, including the small and large cavities with their associated internal and external additional densities. The relative orientation of the two substructures differs between the GPT and GGT folds, due to different conformations of intervening sequences, in particular in the PGGG turn at the end of R2 and in the N-terminal part of R3 that have similar side chain orientations to PSP. In GPT type 2 filaments, the two symmetric inter-protofilament interfaces resemble the asymmetric interface of GGT type 3 filaments, increasing the overall similarity between GGT and GPT filaments (**Extended Data Figure 5**).

AGD is another 4R tauopathy belonging to the spectrum of sporadic FTLD-tau diseases (26–30). A defining feature is the presence of argyrophilic grains. In addition, tau-immunoreactive coiled bodies and pretangles in limbic projection neurons are consistent features. Bush-like astrocytic inclusions can also be present. Argyrophilic grains are rarely the sole pathological finding in cognitively impaired subjects and are most commonly found together with other tau pathologies, especially neurofibrillary tangles (28).

We determined the cryo-EM structures of tau filaments from the nucleus accumbens of two cases of stage 3 AGD with resolutions up to 3.4 Å (**Figure 2**). We observed three different types of filaments with a common protofilament core, the AGD fold, that adopts a four-layered ordered structure comprising residues 273-387 (type 1) or 279-381 (type 2), and resembles the CBD fold (5). Like CBD type I and type II filaments, AGD type 1 filaments consist of a single protofilament and AGD type 2 filaments contain two protofilaments that pack against each other with C2 symmetry. We were unable to solve the structure of AGD type 3 filaments to sufficient resolution for atomic modelling, but low-resolution cross-sections suggest an asymmetric packing arrangement of two protofilaments with the AGD fold. AGD case 1 also had Alzheimer PHFs and CTE type I filaments, whereas AGD case 2 also had Alzheimer PHFs and SFs (**Extended Data Figure 2d**).

**Figure 2:**
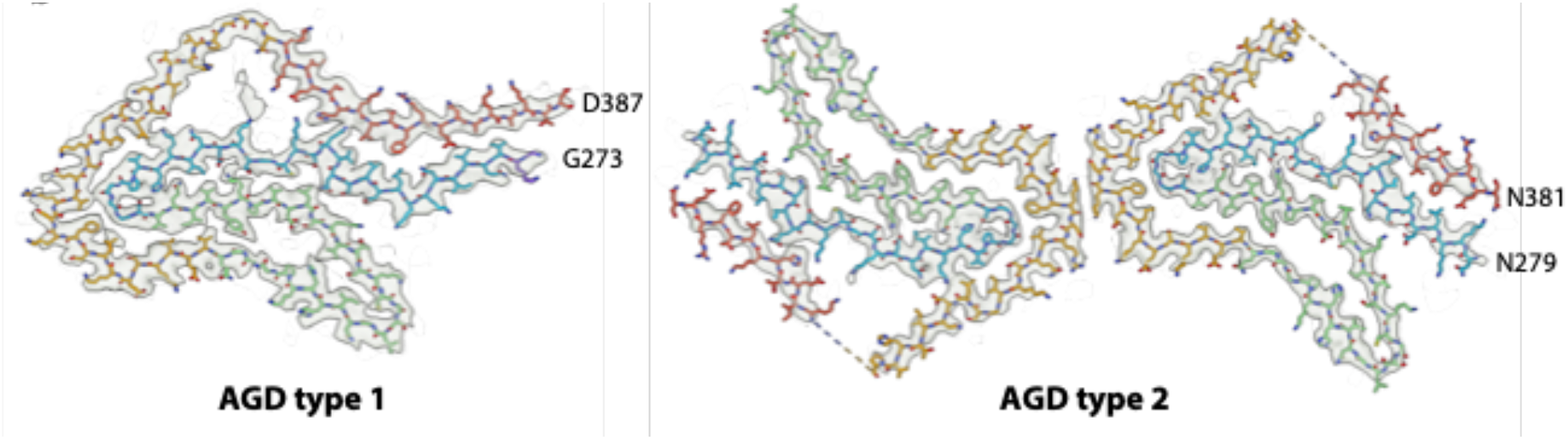
Four-layered 4R tau filament structures. Cryo-EM density map and atomic model for AGD type 1 filaments and AGD type 2 filaments. Colour scheme is as in Figure 1.

A large section of the AGD fold, residues 293-357, adopts the same conformation as in the CBD fold. However, the C-terminal segment (368-386) makes different interactions with R2, and the ordered core of the AGD fold is seven amino acids longer than that of the CBD fold. In addition, compared to the CBD cavity, the AGD internal cavity between R2 and the C-terminal segment is smaller and its net positive charge is less. It also contains a smaller additional density of unknown identity between K294 and K370. In AGD type 2 filaments, the turn at the PGGG motif at the end of R4, as well as the first and last six residues that form the ordered core of the AGD type 1 filaments, are disordered and the termini point in a different direction compared to the AGD type 1 filaments.

Histologically, PSP-RS case 3 had also features of AGD, in particular the presence of argyrophilic grains in the entorhinal cortex (**Extended Data Figure 6**). Consistent with this, besides filaments with the PSP fold in thalamus, we also observed filaments of AGD type 2 and AGD type 3, as well as a minority of Alzheimer PHFs and SFs, in the entorhinal cortex from PSP-RS case 3 (**Extended Data Figure 2e**). By immunoblotting, the pattern of C-terminally truncated tau bands was like that of PSP in thalamus and like that of CBD in entorhinal cortex (**Extended Data Figure 7**). Thus, this case had concomitant PSP and AGD, as has been reported for other cases of PSP (28).

Astroglial tau pathology has been increasingly recognized during aging; these cases have been subsumed under the umbrella term of ARTAG (31), which is characterized by the presence of thorn-shaped and granular/fuzzy astrocytes with 4R tau inclusions (32–34). Like AGD, ARTAG usually coexists with other tau pathologies. Different types of ARTAG can be distinguished in grey and white matter, as well as in subpial, subependymal and perivascular locations (31). We determined the cryo-EM structures of the majority species of tau filaments from the hippocampus of a case of ARTAG (34). Most tau filaments were identical to AGD type 2 and type 3 filaments. Alzheimer PHFs and SFs were also observed.

Unlike the sporadic diseases described above, mutations +3/+16 in intron 10 of *MAPT* give rise to a 4R tauopathy that belongs to the spectrum of inherited FTLD-tau (35–38). It is characterized by abundant filamentous tau inclusions in nerve cells and glial cells in many brain regions. Abundant argyrophilic grains are present in entorhinal cortex and hippocampus from cases with the +3 mutation [multiple system tauopathy with presenile dementia (MSTD)], and some are seen in frontal cortex. Bushy astrocytes are present in hippocampus and frontal cortex. We determined the cryo-EM structures of tau filaments from the frontal cortex of two cases with a +3 and one case with a +16 mutation in intron 10 of *MAPT* (Fig 2A). In all three cases, we only observed the AGD fold.

The differences between the cryo-EM structures of tau filaments from 4R tauopathies are consistent with results from immunoblots of sarkosyl-insoluble tau (**Extended Data Figure 7**) (39–41). In PSP, CBD, GGT, AGD, ARTAG and intron 10 +3/+16 mutation cases, tau bands of 60 and 64 kDa were in evidence, indicating the presence of full-length 4R tau. As reported previously, a strong C-terminal tau band of 33 kDa was also found in PSP and GGT, whereas a strong tau doublet of 37 kDa was seen in CBD, AGD and cases with mutation +16 in intron 10 of *MAPT*. The 37 kDa bands were also characteristic of ARTAG and cases with mutation +3 in intron 10 of *MAPT*.

Whereas all the diseases described above are 4R tauopathies, we also determined structures of tau filaments from two inherited types of dementia that contain mixtures of 3R+4R tau in their neuronal inclusions. FBD and FDD are caused by mutations in the integral membrane protein 2B gene (*ITM2B*) and are characterized by abundant extracellular inclusions of British amyloid (ABri) and Danish amyloid (ADan) that are associated with abundant tau-positive neurofibrillary tangles, neuropil threads and dystrophic neurites (42–45). We determined the cryo-EM structures of tau filaments from the hippocampus of a previously described case of FBD (44) and the temporal cortex of a case of FDD (**Extended Data Figure 2h**). In both cases, we found tau filaments identical to the PHFs that we previously observed for PART and Alzheimer’s disease (1,2,10). The hippocampus from the case of FBD also contained Alzheimer SFs, as well as CTE type I tau filaments, possibly as a result of head trauma resulting from ataxia.

## DISCUSSION

The cryo-EM structures presented here, together with those previously described for Alzheimer’s disease (1,2,10), Pick’s disease (3), CTE (4), CBD (5) and PART (10), provide an overarching perspective on tau filaments that suggests a hierarchical classification of human tauopathies based on their filament folds (**Figure 3**).

**Figure 3:**
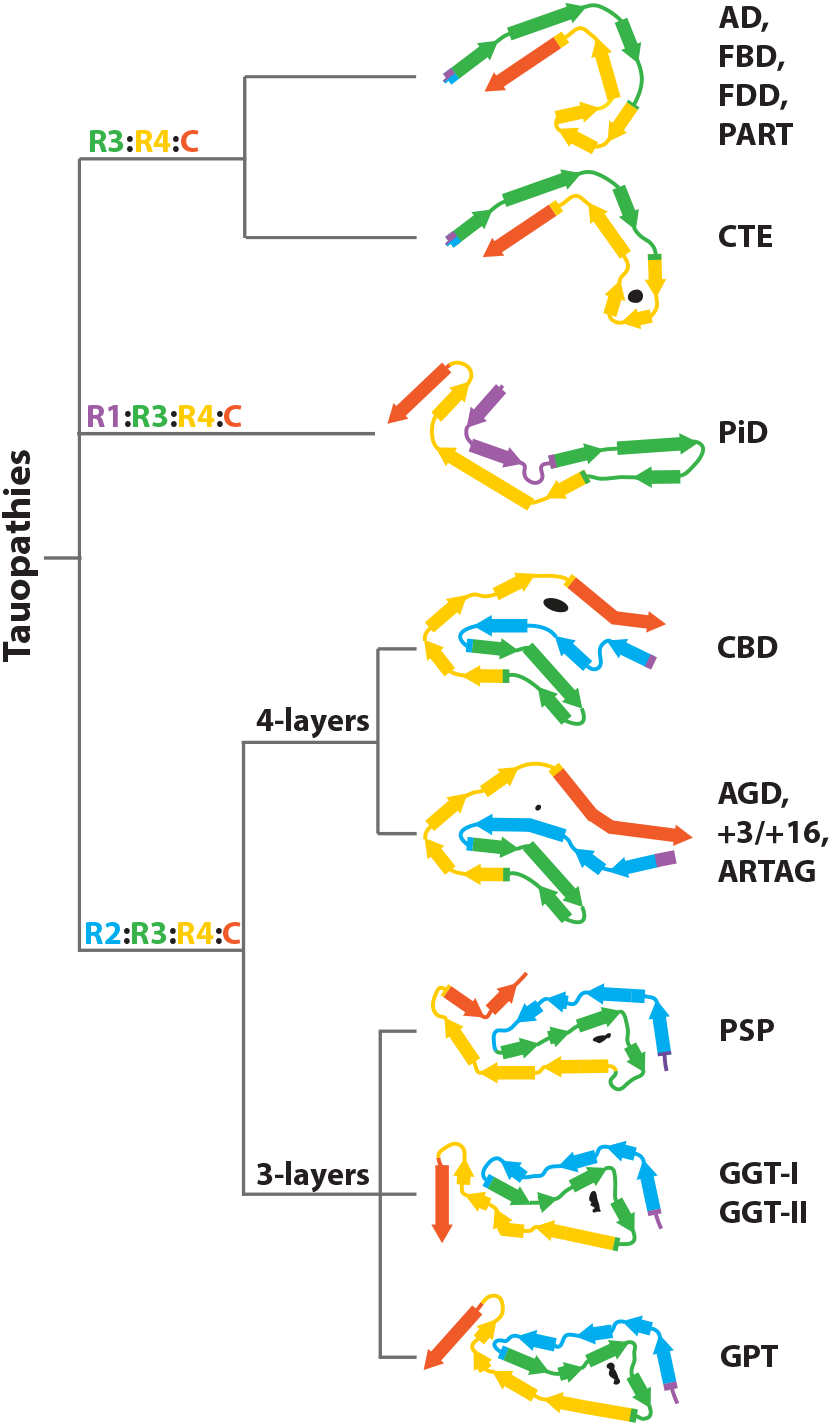
Structure-based classification of tauopathies.

The first level of classification is based on the extent of the ordered cores, and coincides with the different isoform compositions of tau inclusions in the corresponding diseases. All known tau folds from human diseases have a common ordered core region that comprises R3 and R4, as well as 10-13 amino acids of the C-terminal domain. However, these folds differ in their N-terminal extensions. The Alzheimer and CTE folds, which represent 3R+4R tauopathies, extend only one or two residues on the N-terminal side of the common core region. Whereas Alzheimer’s disease, FBD, FDD and PART are distinct diseases that are characterized by different symptoms and/or neuropathology, they all contain tau filaments with the Alzheimer fold. The Pick fold, representing the only 3R tauopathy with known filament structures, comprises more than half of R1. Folds observed for 4R tauopathies comprise all of R2 and one or two residues of R1. As we previously observed for the CBD fold, incorporation of 3R tau isoforms is incompatible with the PSP, GPT, GGT and AGD folds, which agrees with the observation that only 4R tau isoforms assemble in these diseases.

At a second level, 4R tauopathies are divided into two major classes. The PSP, GPT and GGT folds comprise elongated three-layered core regions, whereas the CBD and AGD folds adopt a four-layered topology. The AGD fold is also found in ARTAG, as expected from the presence of granular/fuzzy astrocytes in grey matter in both AGD and ARTAG, and in cases with intron 10 mutations +3/+16 of *MAPT*, consistent with the presence of argyrophilic grains in cases with the +3 mutation (**Extended Data Figure 6**). The division of 4R tauopathies based on three-layered and four-layered structures agrees with observations on posttranslational modifications, which showed similarities between PSP and GGT, and between CBD and cases with mutation +16 in intron 10 of *MAPT* (46).

Assembled tau in AGD has been reported to lack acetylation (47). It is not known if this is also true of ARTAG and cases with +3/+16 intronic mutations. Features resembling AGD have also been described in cases of frontotemporal dementia caused by exon 10 mutations S305I and S305S in *MAPT* (48,49). These exonic and all mutations in intron 10 destabilise the tau exon 10 splicing regulatory element RNA, resulting in a relative overproduction of 4R tau and the formation of 4R tau filaments (50). These filaments may have the AGD fold in common.

At the residue level, the three-layered PSP, GGT and GPT folds are different. Prior to its definition, some cases of GGT were diagnosed as atypical PSP (51). Our observations that the tau filaments from GGT Type I and Type II are distinct from those of PSP, support the definition of GGT as a separate neuropathological entity. Similarly, the observation that the structures of tau filaments from PSP-F case 2 are different from those of PSP and GGT suggests that this individual suffered from a distinct disease, which we named Limbic-predominant Neuronal inclusion body 4R Tauopathy (LNT).

By bridging observations at the molecular level with those at the levels of disease symptoms and neuropathology, our results provide a structural framework for classifying and studying tauopathies.

## Acknowledgements

We thank the patients’ families for donating brain tissues; U. Kuederli, M. Jacobsen, F. Epperson and R.M. Richardson for human brain collection and technical support; E. Gelpi for preparing brain samples from the ARTAG case; E. de Jong, B. van Knippenberg, L. Yu and E. Ioannou for support with the Krios G4 microscope; T. Darling and J. Grimmett for help with high-performance computing; S. Lövestam, T. Nakane, R.A. Crowther, F. Clavaguera, K. Del Tredici, H. Braak and M.G. Spillantini for helpful discussions. We acknowledge Diamond Light Source for access and support of the cryo-EM facilities at the UK’s national Electron Bio-imaging Centre (eBIC) [under proposal BI23268-49], funded by Wellcome Trust, MRC and BBRSC. This study was supported by the MRC-LMB EM facility. W.Z. was supported by a foundation that prefers to remain anonymous. M.G. is an Honorary Professor in the Department of Clinical Neurosciences of the University of Cambridge. This work was supported by the U.K. Medical Research Council (MC_U105184291 to M.G. and MC_UP_A025_1013 to S.H.W.S.), and the EU/EFPIA/Innovative Medicines Initiative [2] Joint Undertaking IMPRiND, project 116060, to M.G.), the Japan Agency for Science and Technology (Crest, JPMJCR18H3), to M.H., the Japan Agency for Medical Research and Development (AMED, JP20dm0207072), to M.H., the Japan Agency for Medical Research and Development (AMED, JP21dk0207045 and JP21ek0109545), to T.I., the U.S. National Institutes of Health (P30-AG010133, UO1-NS110437 and RF1-AG071177), to R.V. and B.G. and the Department of Pathology and Laboratory Medicine, Indiana University School of Medicine. G.G.K. was supported by the Safra Foundation and the Rossy Foundation. T.R. is supported by the National Institute for Health Research Queen Square Biomedical Research Unit in Dementia. T.L. holds an Alzheimer’s Research UK Senior Fellowship. The Queen Square Brain Bank is supported by the Reta Lila Weston Institute for Neurological Studies.

## METHODS

No statistical methods were used to predetermine sample size. The experiments were not randomized and investigators were not blinded to allocation during experiments and outcome assessment.

### Clinical history and neuropathology

We determined the cryo-EM structures of tau filaments from the brains of seven individuals with PSP, one individual with GGT-I, one individual with GGT-II, two individuals with AGD, one individual with ARTAG, two individuals with mutation +3 in intron 10 of *MAPT*, one individual with mutation +16 in intron 10 of *MAPT*, one individual with familial British dementia and one individual with familial Danish dementia. The tau gene was sequenced for all cases. With the exception of the cases with +3 and +16 mutations, the sequences of *MAPT* exons and adjacent introns were wild-type. PSP-RS case 1 was in an 83-year-old man who died with a neuropathologically confirmed diagnosis following a 4-year history of falls, bradykinesia, cognitive impairment and supranuclear gaze palsy. Microscopic features showed degeneration of pallidum, tegmentum and dentate nucleus of the cerebellum. Abundant 4R tau-positive tangles, neuropil threads, tufted astrocytes and coiled bodies were in evidence. PSP-RS case 2 was in a 70-year-old man who died with a neuropathologically confirmed diagnosis following a 7-year history of postural instability, gait disturbance, cognitive impairment and supranuclear gaze palsy. Magnetic resonance imaging (MRI) of the brain showed the hummingbird sign, probably reflecting a severe midbrain atrophy. PSP-RS case 3 was in a 74-year-old man who died following a 5-year history of postural instability, dysphagia and cognitive dysfunction. Neuronal loss and gliosis, as well as abundant 4R tau-immunoreactive neuronal and glial tau inclusions were present in thalamus, subthalamic nucleus, cerebellum, midbrain, pons and medulla; 4R tau-immunoreactive argyrophilic grains were in evidence in entorhinal cortex and hippocampus. The neuropathological diagnosis was PSP + AGD. PSP-F case 1 was in a 62-year-old man who died with a neuropathologically confirmed diagnosis following an 11-year history of behavioural-variant FTD, falls, bradykinesia and dysphagia. Abundant neuronal and glial 4R tau inclusions were present in cortical and, especially, subcortical areas. Basal ganglia, thalamus and brainstem were severely affected. PSP-F case 2 was in a 66-year-old woman who died with FTD and parkinsonism. Abundant neuronal and glial 4R tau inclusions were present in cortical and subcortical areas. In limbic areas and other brain regions, abundant spherical, 4R tau-immunoreactive, basophilic and Gallyas silver-positive neuronal inclusions were present, together with globular glial tau inclusions. PSP-P was in a 63-year-old man who died with a neuropathologically confirmed diagnosis following a 9-year history of akinetic-rigid parkinsonism, falls and cognitive impairment. Brain MRI showed progressive frontal atrophy and mild midbrain atrophy. Neuronal loss was more marked in the substantia nigra than in the pallidum, tegmentum and dentate nucleus of the cerebellum. Abundant neuronal and glial 4R tau pathology was widespread. PSP-CBS was in an 88-year-old woman with a neuropathologically confirmed diagnosis following a 9-year history of cerebellar ataxia, corticobasal syndrome and cognitive impairment. Abundant neuronal and glial 4R tau inclusions were present in cerebral cortex, from the precentral gyrus to the operculum. GGT-I was in a 77-year-old woman with a neuropathologically confirmed diagnosis following a 3-year history of falls, bradykinesia and cognitive impairment. The clinical diagnosis was behavioural-variant FTD with an atypical parkinsonian disorder (19). Neuropathologically, severe anterior frontal and temporal cortical nerve cell loss and severe tau deposition in cerebral cortex and subcortical white matter were in evidence. Abundant neuronal, as well as globular oligodendroglial and astrocytic 4R tau inclusions, were present. GGT-II was in a 76-year-old woman with a neuropathologically confirmed diagnosis following a 5-year history of asymmetric pyramidal signs, diffuse muscle atrophy and dysarthria (20). Severe neuronal loss was present in the motor cortex, with an almost complete loss of Betz cells. Abundant 4R tau neuronal cytoplasmic inclusions, globular oligodendroglial inclusions, coiled bodies and neuropil threads were in evidence. Only a small number of globular astroglial inclusions was seen in the affected cerebral cortex. GGT-III was in a 62-year-old man with a neuropathologically confirmed diagnosis following a 4-year history of parkinsonism, motor signs and dementia (21). Loss of anterior horn cells and degeneration of the corticospinal tract were in evidence. Abundant 4R tau neuronal and globular oligodendroglial and astroglial inclusions were present. Tau filament structures were determined from the nucleus accumbens of two cases of AGD with a predominantly neuronal 4R tau pathology. The first case was in a 90-year-old man with a 6-year history of memory loss and behavioural changes. By imaging, diffuse atrophy of the medial temporal lobe was observed. At autopsy, a chronic subdural hematoma was present on the right side, supporting the possibility of a previous head injury and consistent with the presence of CTE type I tau filaments in the nucleus accumbens. However, there was no history of head trauma. Nerve cell loss and numerous tau-immunoreactive argyrophilic grains were observed in entorhinal cortex, hippocampus and amygdala. Some tau-immunoreactive coiled bodies and bush-like astrocytes were also present. The second case of AGD was in an 85-year-old man with a 3-year history of falls and memory loss. At autopsy, atrophy of hippocampus, amygdaloid nucleus and parahippocampal gyrus was present. Abundant 4R tau-immunoreactive argyrophilic grains were in evidence, with some coiled bodies and bush-like astrocytes. ARTAG was identified in an 85-year-old woman with a 1-year history of cancer and depression (34). At autopsy, she had prominent ARTAG (subpial, subependymal, grey matter, white matter and perivascular). A heterozygous mutation at position +3 in intron 10 of *MAPT* was identified in two patients with dementia and a family history of MSTD. The first individual was a 54-year-old woman who died following a 7-year history of disinhibition, anxiety and cognitive impairment. The second individual with a +3 mutation in intron 10 of *MAPT* was a 63-year-old woman who died following an 8-year history of dementia with a severe amnestic component. In both individuals, nerve cell loss and gliosis were severe in neocortex, amygdala, entorhinal cortex, hippocampus, basal ganglia, subthalamic nucleus, midbrain and brainstem. Widespread silver-positive neuronal, oligodendroglial and astrocytic 4R tau inclusions were in evidence. A mutation at position +16 in intron 10 of *MAPT* was identified in a 53-year-old man with a 7-year history of behavioural-variant FTD. This subject had a family history of FTD, consistent with dominant inheritance. Abundant 4R tau neuronal and glial cell inclusions were in evidence. Some glial cell inclusions resembled astrocytic plaques. FBD (T to A mutation, removing the stop codon in *ITM2B*) was in a 68-year-old woman with an 11-year history of ataxia and progressive cognitive decline. Her speech became slurred and she experienced problems with swallowing. A brother and two uncles had similar symptoms. As reported previously [case 5 in (44)], neuropathological findings included degeneration of the hemispheric white matter, parenchymal and blood vessel deposits of ABri amyloid, as well as widespread neurofibrillary pathology. This patient had a serious bicycle accident, probably as a result of ataxia, which may explain the presence of some CTE type I filaments. FDD (decamer duplication between codons 265 and 266 of *ITM2B*) was in a 52-year-old patient from generation V of a previously described kindred (45). He developed cataracts aged 21 and had severe hearing loss, nystagmus and ataxia aged 38. Neuropathological examination showed similar findings as in FBD, with a difference being that ADan, which differs from ABri amyloid in its 12 C-terminal amino acids, formed mostly thioflavin S-negative deposits.

### Whole-exome sequencing

Target enrichment made use of the SureSelectTX human all-exon library (V6, 58 mega base pairs; Agilent) and high-throughput sequencing was carried out using a HiSeq4,000 (2×75-base-pair paired-end configuration; Illumina). Bioinformatics analyses were performed as described (52)

### Extraction of tau filaments

For cryo-EM, sarkosyl-insoluble material was extracted from frontal cortex (PSP-RS1, PSP-RS2, PSP-P, PSP-CBS, GGT-I, GGT-II, GGT-III, +3 case 1, +3 case 2, +16), temporal cortex (PSP-F2, FDD), entorhinal cortex (PSP-RS3), hippocampus (FBD, ARTAG), putamen (PSP-F1), thalamus (PSP-RS3) and nucleus accumbens (AGD cases 1 and 2), essentially as described (40). Briefly, tissues were homogenized in 20 volumes (v/w) extraction buffer consisting of 10 mM Tris-HCl, pH 7.5, 0.8 M NaCl, 10% sucrose and 1 mM EGTA. Homogenates were brought to 2% sarkosyl and incubated for 30 min. at 37°C. For PSP-RS case 1, PSP-RS case 2, PSP-CBS, PSP-P, GGT-II, GGT-III, AGD case 1, AGD case 2, +3 case 1, +3 case 2, +16 case, following a 10 min. centrifugation at 20,000 g, the supernatants were spun at 100,000 g for 20 min. For the rest of the cases, the supernatants from a 10 min. centrifugation at 7,000 g were spun at 100,000 g for 60 min. The pellets were resuspended in 700 μl/g extraction buffer and centrifuged at 9,500 g for 10 min. For PSP-RS case 1, PSP-RS case 2, PSP-CBS, PSP-P, GGT-II, GGT-III, AGD case 1, AGD case 2, and +16 case, the supernatants were diluted 3-fold in 50 mM Tris-HCl, pH 7.5, containing 0.15 M NaCl, 10% sucrose and 0.2% sarkosyl and spun at 166,000 g for 30 min. For the other cases, the supernatants were spun at 100,000 g for 60 min., the pellets resuspended in 700 μl/g extraction buffer and centrifuged at 9,800 g. The supernatants were then spun at 100,000 g for 60 min. Sarkosyl-insoluble pellets were resuspended in 25 μl/g of 20 mM Tris-HCl, pH 7.4, 100 mM NaCl and used for cryo-EM.

### Immunoblotting, histology and silver staining

Immunoblotting was carried out as described (40). Samples were resolved using 4-20% Tris-glycine gels (Novex) and antibody T46 was used at 1:2,000. Histology and immunohistochemistry were carried out as described (53). Brain sections were 8 μm thick and were counterstained with haematoxylin. Primary antibodies were: RD3 (1:1,000); RD4 (1:1,000); AT8 (1:300). Sections were silver-impreganated using the method of Gallyas-Braak (54).

### Electron cryo-microscopy

Extracted tau filaments were centrifuged at 3,000 g for 60 s, before being applied to glow-discharged holey carbon grids (Quantifoil Au R1.2/1.3, 300 mesh) and plunge-frozen in liquid ethane using a Thermo Fisher Vitrobot Mark IV. Images for PSP-F case 2 were acquired on a 300 keV Thermo Fisher Titan Krios G4 microscope, described previously (25), which was equipped with a cold field-emission gun, a Selectrix X energy filter, and a Falcon-IV detector. The energy filter was operated with a slit width of 10 eV. Images were recorded at a total dose of 50 e^-^/Å^2^ using aberration-free image shift (AFIS) as incorporated within the Thermo Fisher EPU software at a throughput of 300 images/h in EER format (55). All other data sets were acquired on Thermo Fisher Titan Krios microscopes with either a Gatan K2 or K3 detector in counting mode, using a GIF-quantum energy filter (Gatan) with a slit width of 20 eV to remove inelastically scattered electrons. Further details are given in Extended Data Table 1.

### Helical reconstruction

Movie frames were gain-corrected, aligned, dose-weighted and then summed into a single micrograph using RELION’s own motion correction program (56). The micrographs were used to estimate the contrast transfer function (CTF) using CTFFIND-4.1 (57). All subsequent image-processing steps were performed using helical reconstruction methods in RELION (58,59). The filaments from PSP-F2, FBD and FDD were picked by crYOLO (60); the rest of the filaments were selected manually in the micrographs. For +3 and +16 cases, filaments of different types were picked separately; for the other data sets different filament types were separated based on the appearance of 2D class averages. For all data sets, reference-free 2D classification was performed to select suitable segments for further processing. Initial 3D reference models were generated *de novo* from the 2D class averages using an estimated rise of 4.75 Å and helical twists according to the observed cross-over distances of the filaments in the micrographs (61) for all data sets, except those from thalamus of PSP-RS case 3, PSP-P, PSP-F case 1, GGT-II, AGD case 2 and ARTAG. Combinations of 3D autorefinements and 3D classifications were then used to select the best segments for each structure. For all data sets, Bayesian polishing (56) and CTF refinement (62) were performed to further increase the resolution of the reconstructions. For PSP-F case 2, temporal drift in the beam tilt was corrected by grouping every 8,000 consecutively collected segments into separate optics groups before CTF refinement. Final reconstructions were sharpened using the standard postprocessing procedures in RELION, and overall final resolutions were estimated from Fourier shell correlations at 0.143 between the two independently refined halfmaps, using phase-randomisation to correct for convolution effects of a generous, soft-edged solvent mask (63). Local resolution estimates were obtained using the same phase-randomisation procedure, but with a soft spherical mask that was moved over the entire map. Further details of data acquisition and processing are given in **Extended Data Tables 1-4**.

### Model building and refinement

Where multiple structures from different cases were obtained, atomic models were only built and refined in the best available maps. For the PSP structure, this was PSP-RS case 1; for the GGT structures, this was GGT-I; for the GPT structures this was PSP-F case 2; for the AGD type 1 filament, this was AGD case 1; for the AGD type 2 filament, this was +16. Atomic models were built manually using COOT (64). Side chain clashes were detected using MOLPROBITY (65) and corrected by iterative cycles of real-space refinement in COOT and Fourier-space refinement in REFMAC (66) and/or real-space refinement in PHENIX (67). For each refined structure, separate model refinements were performed against a single half-map, and the resulting model was compared to the other half-map to confirm the absence of overfitting (**Extended Data Figure 3**). Statistics for the final models are shown in **Extended Data Table 1**.

## Ethical review processes and informed consent

The studies carried out at Tokyo Metropolitan Institute of Medical Science, Indiana University, UCL Queen Square Institute of Neurology, and the University of Toronto were approved through the ethical review processes at each Institution. Informed consent was obtained from the patients’ next of kin.

## Data availability

Cryo-EM maps have been deposited in the Electron Microscopy Data Bank (EMDB) under accession numbers EMD-xxxx. Corresponding refined atomic models have been deposited in the Protein Data Bank (PDB) under accession numbers XXXX. Raw cryo-EM data for data sets XXX have been deposited at EMPIAR under accession numbers XXXX. Any other relevant data are available from the corresponding authors upon reasonable request.

## Extended Data Figures

**Extended Data Figure 1:**
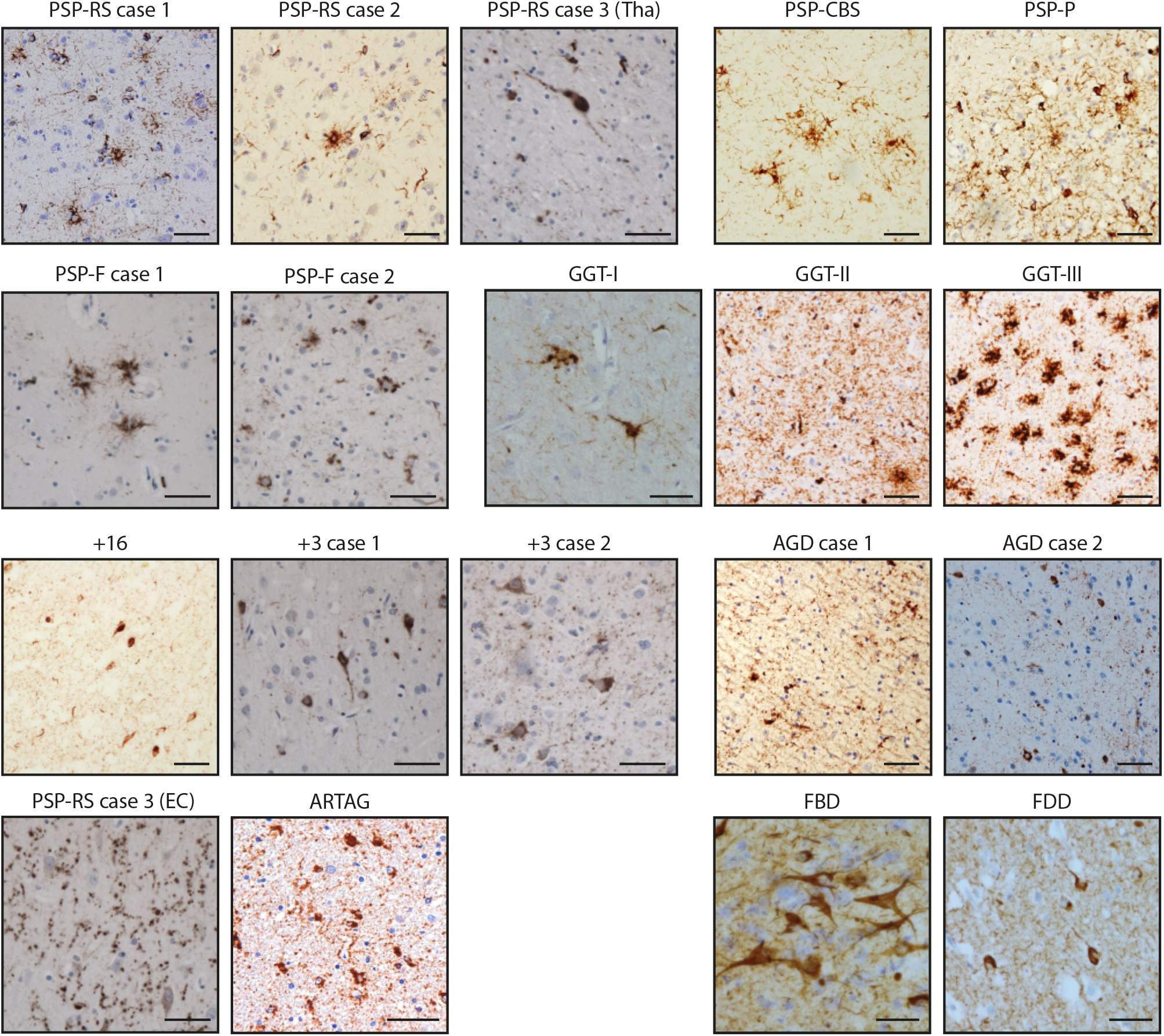
Tau immunohistochemistry. Tau staining of the brain regions used for cryo-EM structure determination (see Methods), using antibody AT8 (pS202/pT205 tau). Scale bars are 50 μm, except for GGT Type I, FBD and FDD, where they are 25 μm. For PSP-RS case 3, both the thalamus (Tha) and the entorhinal cortex (EC) are shown.

**Extended Data Figure 2:**
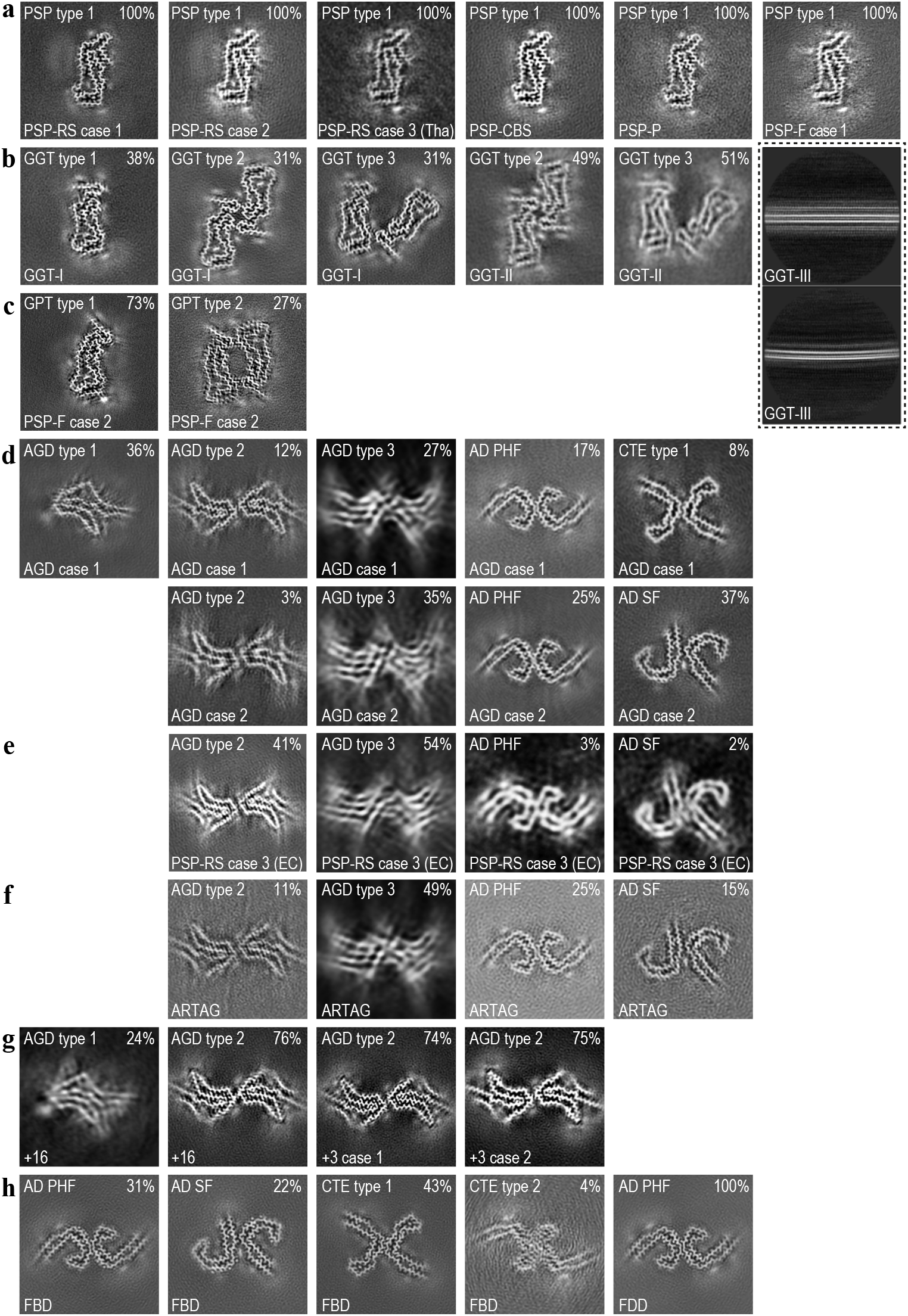
Cryo-EM reconstructions. For each map, a sum of the reconstructed density for several central XY-slices, corresponding to ~4.7 Å, is shown. The disease cases are referenced at the bottom of each image; the filament type at the top left; and the percentage of that filament type among the tau filaments in the data set at the top right. The inset with dashed lines shows 2D class average images of GGT-III filaments without apparent twist.

**Extended Data Figure 3:**
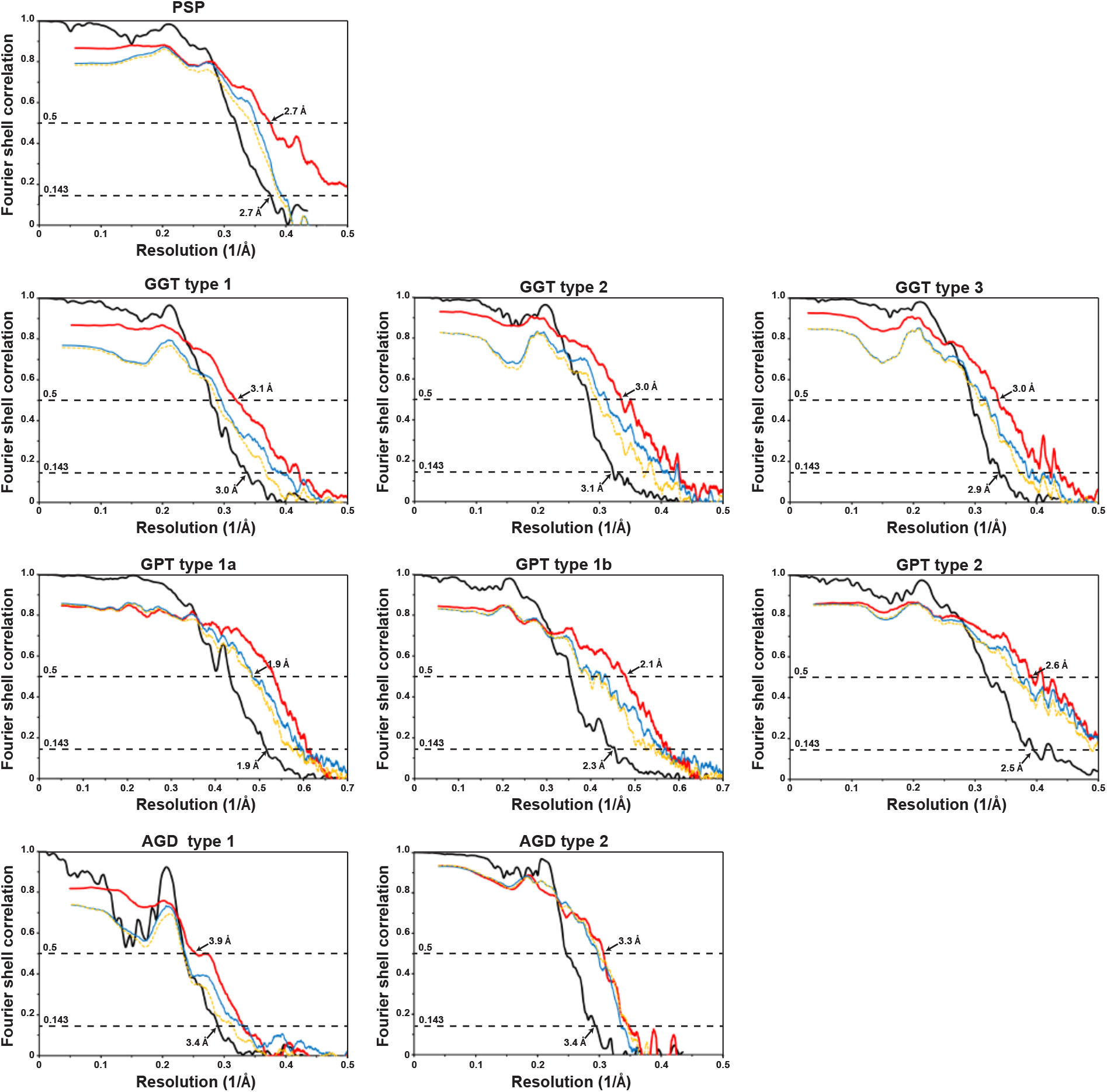
Cryo-EM resolution estimates. Fourier Shell Correlation (FSC) curves for cryo-EM maps and atomic structures of PSP filaments (from PSP-RS case 1); GGT filament types 1-3 (from GGT-I); GPT filament types 1a, 1b and 2 (from PSP-F case 2); and AGD filament types 1 and 2 (from AGD case 1 and the +16 case, repectively). FSC curves are shown for two independently refined cryo-EM half-maps (black); for the final refined atomic model against the final cryo-EM map (red); for the atomic model refined in the first halfmap against that half-map (blue); and for the refined atomic model in the first halfmap against the other half-map (yellow).

**Extended Data Figure 4:**
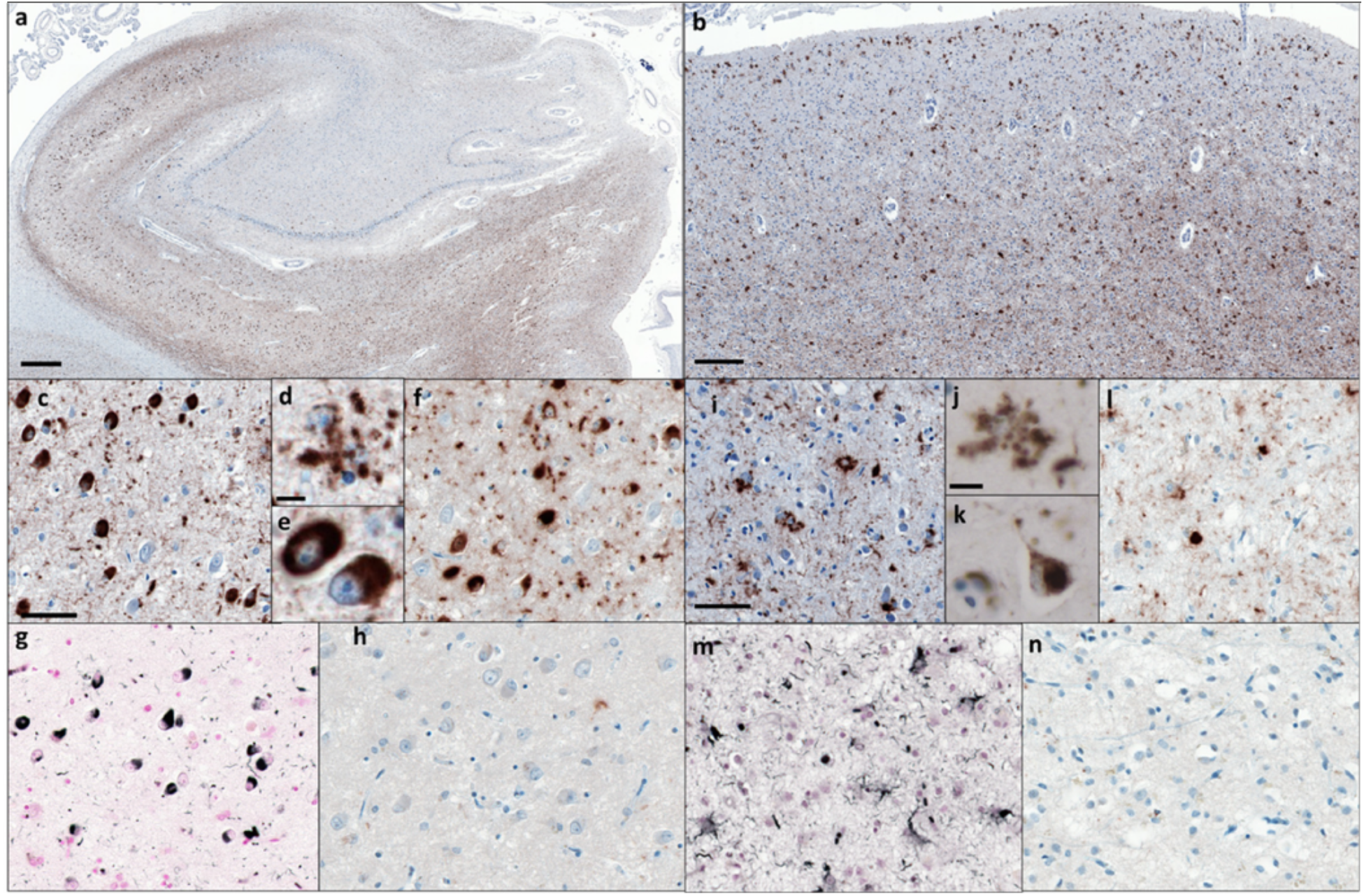
Tau pathology in Limbic-predominant Neuronal inclusion body 4R Tauopathy (LNT, PSP-F case 2) **a)** Low power view of hippocampus stained with antibody AT8 (pS202/pT205 tau). **b)** Low power view of frontal cortex stained with AT8. **c)** Higher power view of hippocampus stained with AT8. **d)** AT8-positive globular astrocyte in hippocampus. **e)** AT8-positive neurons in hippocampus. **f)** Hippocampus stained with antibody RD4 (specific for 4R tau). **g)** Gallyas-Braak silver-positive neurons and glial cells in hippocampus. **h)** Hippocampus stained with antibody RD3 (specific for 3R tau). **i)** Higher power view of frontal cortex stained with AT8. **j)** AT8-positive globular astrocyte in frontal cortex. **k)** AT8-positive neruron in frontal cortex. **l)** Frontal cortex stained with RD4. **m)** Gallyas-Braak silver-positive neurons and glial cells in frontal cortex. **n)** Frontal cortex stained with RD3. Scale bars: 400 μm (in a); 200 μm (in b), 50 μm (in c, f, g, h, i, l, m, n); 10 μm (in d, e, j, k).

**Extended Data Figure 5:**
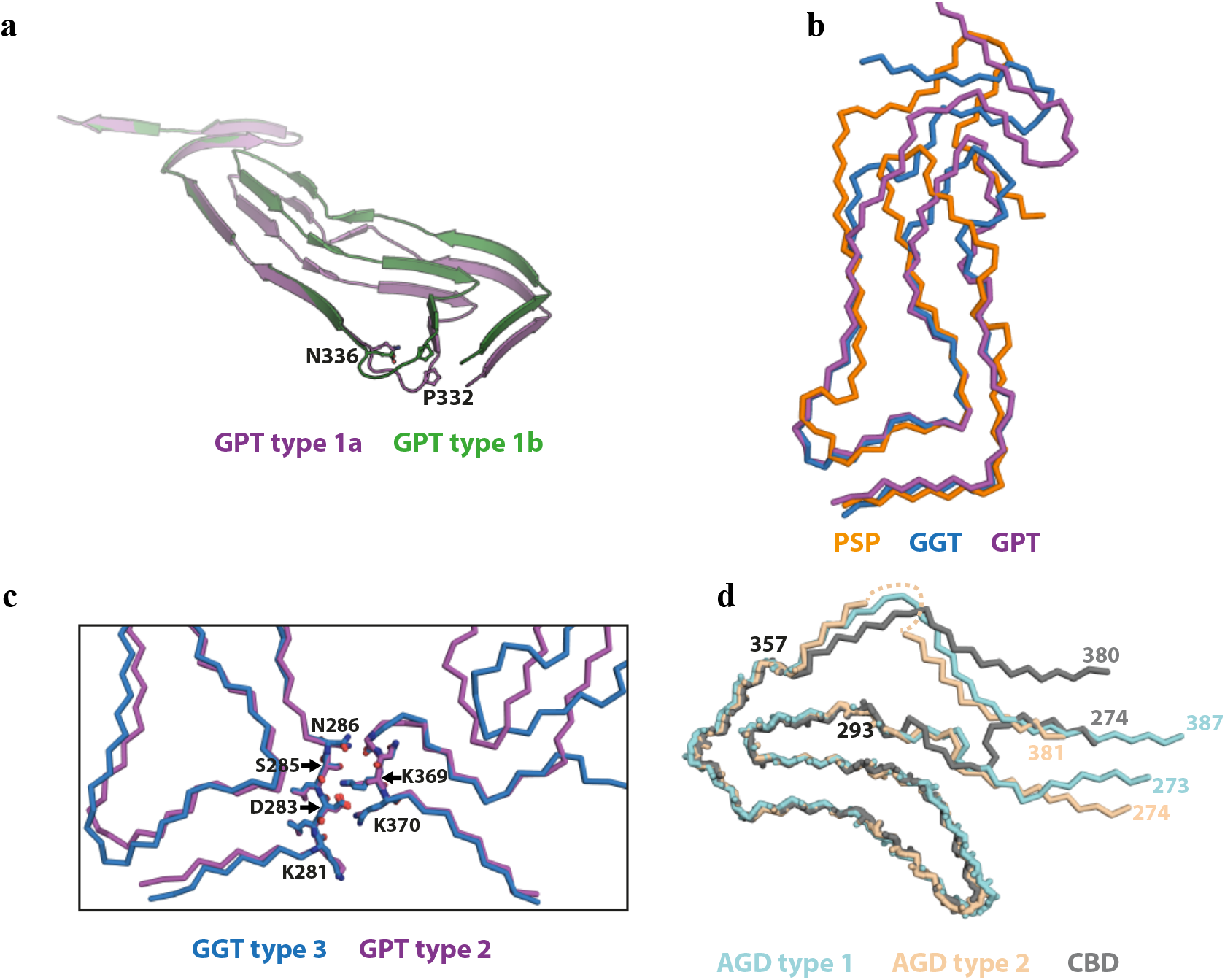
Structural comparisons. **a)** Two different main-chain conformations for GPT type 1 filaments (type 1a in purple; type 1b in green). **b)** Comparison of the PSP (orange), GGT (blue) and GPT (purple) folds. **c)** Comparison of the inter-protofilament interface of GGT type 3 and GPT type 2 filaments. **d)** Comparison of the AGD type 1 (light blue), AGD type 2 (pink) and CBD (gray) folds.

**Extended Data Figure 6:**
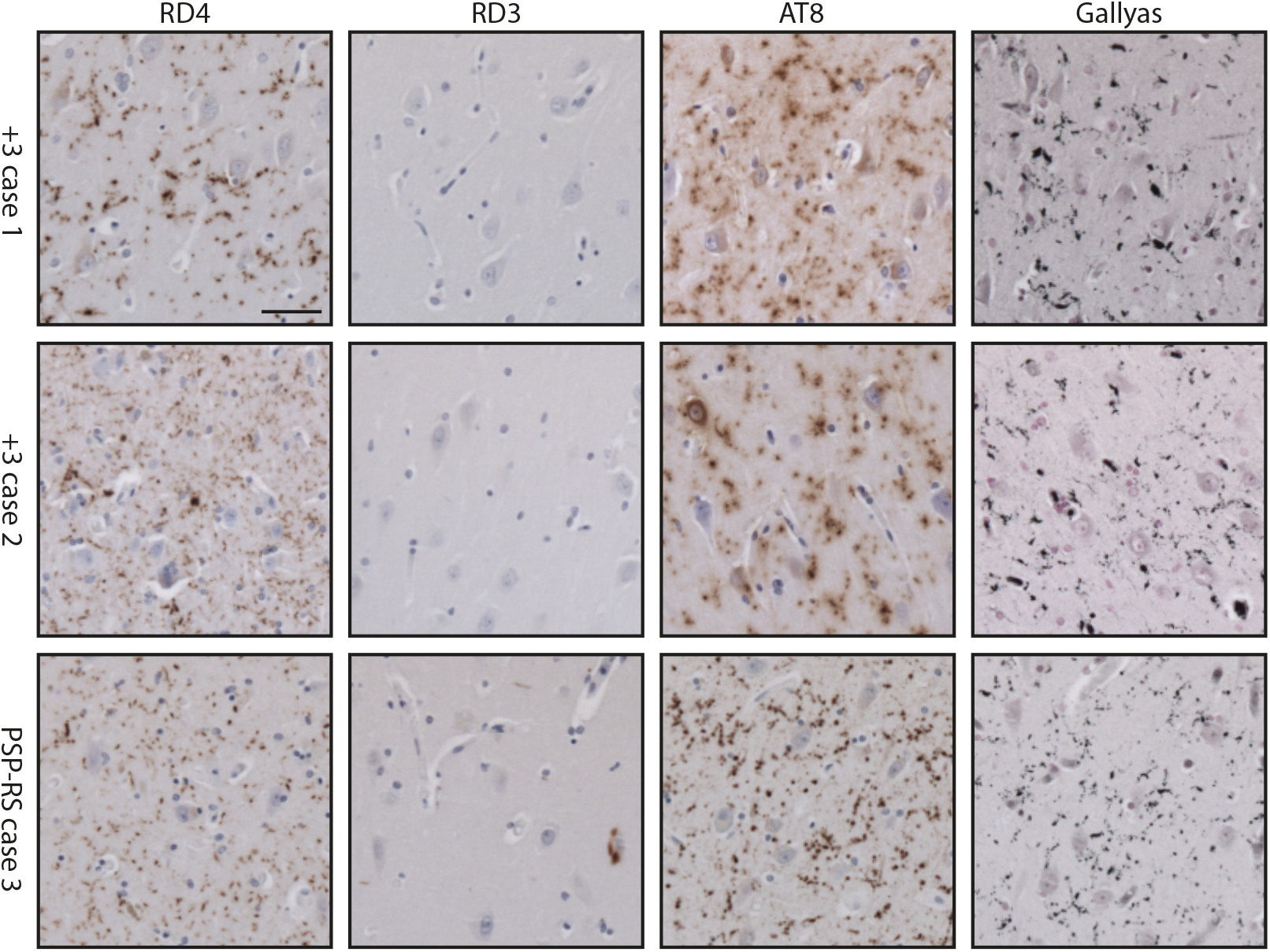
Argyrophilic grains in the entorhinal cortex. Tau staining with antibodies RD4 (4R tau), RD3 (3R tau), AT8 (pS202/pT205 tau), as well as Gallyas-Braak silver, of the entorhinal cortex from cases 1 and 2 of MSTD (+3 intron 10 mutation in *MAPT*) and from case 3 of PSP-RS. Scale bar, 50 μm.

**Extended Data Figure 7:**
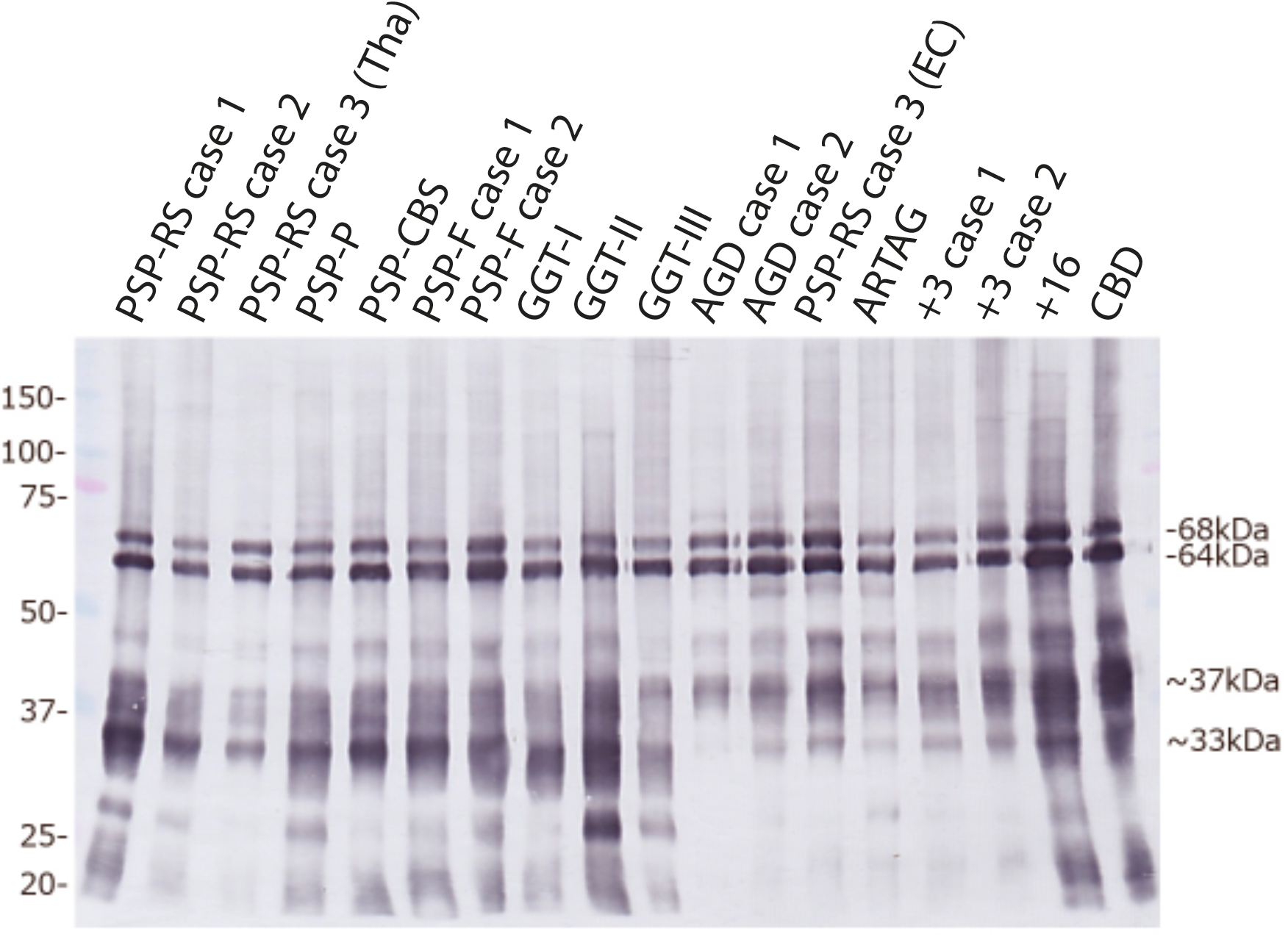
Immunoblot analysis of 4R tauopathies. Hyperphosphorylated full-length tau (64 and 68 kDa) and C-terminal fragments (33 kDa and 37 kDa) were detected in sarkosyl-insoluble fractions from the brain regions used for cryo-EM by anti-tau antibody T46. A prominent 33 kDa band was characteristic of PSP and GGT; strong 37 kDa bands were in evidence in AGD, ARTAG, cases with intron 10 mutations in *MAPT* (+3 and +16) and in CBD. PSP-RS case 3 had a strong 33 kDa band in thalamus (Tha) and strong 37 kDa bands in entorhinal cortex (EC), consistent with AGD co-pathology.

