## Extended Data Tables 1-4 for "Structure-based Classification of Tauopathies"

### Extended Data Table 1 | Cryo-EM data collection, refinement and validation statistics

| | PSP-RS case 1 | GGT-I case 1 | | | PSP-F case 2 | | | +16 case 1 | |
|---|---|---|---|---|---|---|---|---|---|
| **Data collection and processing** | | | | | | | | | |
| Electron Gun | XFEG | XFEG | | | CFEG | | | XFEG | |
| Detector | K2 | K2 | | | Falcon4 | | | K2 | |
| Magnification | 105k | 105k | | | 165k | | | 105k | |
| Voltage (kV) | 300 | 300 | | | 300 | | | 300 | |
| Electron exposure (e–/Å$^2$) | 40 | 50 | | | 50 | | | 54 | |
| Defocus range (μm) | 1.7-2.2 | 1.5-2.5 | | | 0.8-2.0 | | | 1.7-2.8 | |
| Pixel size (Å) | 1.15 | 1.15 | | | 0.727 | | | 1.15 | |
| Initial particle images (no.) | 155,030 | 408,018 | | | 630,640 | | | 375,898 | |
| | | type 1 | type 2 | type 3 | type 1a | type 1b | type 2 | type 1 | type 2 |
| Symmetry imposed | C1 | C1 | C1 | C1 | C1 | C1 | C1 | C1 | C2 |
| Final particle images (no.) | 17,316 | 28,058 | 42,381 | 84,438 | 53,363 | 15,096 | 16,813 | 14,582 | 52,134 |
| Map resolution (Å; FSC=0.143) | 2.7 | 3.0 | 3.1 | 2.9 | 1.9 | 2.3 | 2.5 | 5.7 | 3.4 |
| Helical rise (Å) | 4.78 | 4.75 | 2.37 | 4.75 | 4.78 | 4.78 | 2.39 | 4.80 | 4.80 |
| Helical twist ($^o$) | -0.86 | -0.75 | 179.65 | -0.73 | -0.36 | -0.39 | 179.82 | -0.36 | -0.40 |
| | PSP-RS case 1 type 1 | GGT-I case 1 type 1 | GGT-I case 1 type 2 | GGT-I case 1 type 3 | PSP-F case 2 type 1a | PSP-F case 2 type 1b | PSP-F case 2 type 2 | AGD case 1 type 1 | +16 case 1 type 2 |
| **Refinement** | | | | | | | | | |
| Initial model used (PDB code) | - | - | - | - | - | - | - | 6TJX | 6TJX |
| Model resolution (Å) | 2.7 | 3.1 | 3.0 | 3.0 | 1.9 | 2.1 | 2.6 | 3.9 | 3.3 |
| FSC threshold | | | | | | | | | |
| Map sharpening *B* factor (Å$^2$) | -24.89 | -20.19 | -34.53 | -29.03 | -18.89 | -20.16 | -23.15 | -35.52 | -35.83 |
| Model composition | | | | | | | | | |
| Non-hydrogen atoms | 4145 | 4060 | 8210 | 8120 | 4166 | 4060 | 8120 | 4340 | 4854 |
| Protein residues | 550 | 540 | 1090 | 1080 | 540 | 540 | 1080 | 575 | 630 |
| Waters | - | - | - | - | 76 | - | - | - | - |
| *B* factors (Å$^2$) | | | | | | | | | |
| Protein | 23.36 | 68.49 | 36.29 | 70.77 | 25.15 | 35.28 | 50.31 | 71.32 | 107.39 |
| Waters | - | - | - | - | 30.00 | - | - | - | - |
| R.m.s. deviations | | | | | | | | | |
| Bond lengths (Å) | 0.004 | 0.011 | 0.010 | 0.010 | 0.005 | 0.004 | 0.003 | 0.003 | 0.009 |
| Bond angles (°) | 0.742 | 1.197 | 0.914 | 1.094 | 0.779 | 0.824 | 0.726 | 0.641 | 1.298 |
| Validation | | | | | | | | | |
| MolProbity score | 1.19 | 1.61 | 1.68 | 1.54 | 1.53 | 1.48 | 1.54 | 1.60 | 1.65 |
| Clashscore | 2.60 | 3.97 | 6.44 | 3.43 | 5.23 | 4.45 | 3.61 | 8.90 | 4.83 |
| Poor rotamers (%) | 0.00 | 0.00 | 0.00 | 0.00 | 0.00 | 0.00 | 0.00 | 0.00 | 0.00 |
| Ramachandran plot | | | | | | | | | |
| Favored (%) | 97.22 | 93.40 | 95.33 | 93.87 | 96.23 | 96.23 | 94.34 | 97.35 | 94.06 |
| Allowed (%) | 2.78 | 6.60 | 4.67 | 6.13 | 3.77 | 3.77 | 5.66 | 2.65 | 5.94 |
| Disallowed (%) | 0.00 | 0.00 | 0.00 | 0.00 | 0.00 | 0.00 | 0.00 | 0.00 | 0.00 |
| PDB | | | | | | | | | |
| EMDB | | | | | | | | | |
| EMPIAR | | | | | | | | | |

### Extended Data Table 2 | Cryo-EM data collection of other PSP and GGT cases

| | PSP-RS case 2 | PSP-RS case 3 (T) | PSP-CBS case 1 | PSP-P case 1 | PSP-F 1 case 1 | GGT-II case 1 | GGT-III case 1 |
|---|---|---|---|---|---|---|---|
| **Data collection and processing** | | | | | | | |
| Electron Gun | XFEG | XFEG | XFEG | XFEG | XFEG | XFEG | XFEG |
| Detector | K2 | K2 | K2 | K2 | K2 | K2 | K2 |
| Magnification | 105k | 105k | 105k | 105k | 105k | 105k | 105k |
| Voltage (kV) | 300 | 300 | 300 | 300 | 300 | 300 | 300 |
| Electron exposure (e–/Å$^2$) | 40 | 39 | 51 | 54 | 40 | 47 | 52 |
| Defocus range (µm) | 1.4-2.5 | 1.5-2.5 | 1.0-3.2 | 1.0-2.7 | 1.2-2.7 | 1.0-2.5 | 1.2-2.5 |
| Pixel size (Å) | 1.15 | 1.15 | 1.15 | 1.15 | 1.065 | 1.15 | 0.83 |
| Symmetry imposed | C1 | C1 | C1 | C1 | C1 | Type 2: C1<br>Type 3: C1 | |
| Initial particle images (no.) | 44,221 | 88,114 | 80,637 | 66,597 | 37,847 | 97,787 | |
| Final particle images (no.) | 23,640 | 3,006 | 26,073 | 3,782 | 25,178 | Type 2: 23,377<br>Type 3: 17,671 | |
| Map resolution (Å; FSC=0.143) | 3.6 | 3.7 | 3.5 | 3.4 | 4.0 | Type 2: 4.0<br>Type 3: 4.9 | |
| Helical rise (Å) | 4.78 | 4.78 | 4.80 | 4.77 | 4.84 | Type 2: 4.76<br>Type 3: 4.79 | |
| Helical twist (°) | -0.81 | -0.83 | -0.88 | -0.88 | -0.86 | Type 2: -0.68<br>Type 3: -0.70 | |

### Extended Data Table 3 | Cryo-EM data collection of AGD, ARTAG, +3 cases, and PSP-RS case 3 (EC)

| | AGD case 1 | AGD case 2 | PSP-RS case 3 (EC) | ARTAG case 1 | +3 case 1 | +3 case 2 |
|---|---|---|---|---|---|---|
| **Data collection and processing** | | | | | | |
| Electron Gun | XFEG | XFEG | XFEG | XFEG | XFEG | XFEG |
| Detector | K3 | K3 | K3 | K3 | K2 | K2 |
| Magnification | 64k | 64k | 64k | 64k | 81k | 105k |
| Voltage (kV) | 300 | 300 | 300 | 300 | 300 | 300 |
| Electron exposure (e–/Å$^2$) | 31 | 30 | 37 | 40 | 52 | 53 |
| Defocus range (µm) | 1.0-3.0 | 1.5-2.5 | 1.5-2.5 | 1.2-2.8 | 1.7-2.8 | 1.7-2.8 |
| Pixel size (Å) | 1.17 | 1.17 | 1.17 | 1.17 | 1.06 | 1.15 |
| Symmetry imposed | Type 1: C1<br>Type 2: C2<br>Type 3: C1<br>PHF:C1<br>CTE Type 1:C1 | Type 2: C2<br>Type 3: C1<br>PHF: C1<br>SF: C1 | AGD Type 2: C2<br>AGD Type 3: C1<br>PHF: C1<br>SF: C1 | AGD Type 2: C2<br>AGD Type 3: C1<br>PHF: C1<br>SF:C1 | AGD Type 2: C2 | AGD Type 2: C2 |
| Initial particle images (no.) | 414,341 | 495,315 | 166,987 | 389,113 | 884,933 | 809,408 |
| Final particle images (no.) | Type 1: 9,305<br>Type 2: 53,872<br>Type 3: 82,054<br>PHF: 57,725<br>CTE Type 1: 28,549 | Type 2: 40,827<br>Type 3: 42,139<br>PHF: 75,614<br>SF: 110,573 | AGD Type 2: 17,238<br>AGD Type 3: 48,964<br>PHF: 2,396<br>SF: 2,230 | AGD Type 2: 30,626<br>AGD Type 3: 137,217<br>PHF: 68,972<br>SF: 41,151 | AGD Type 2: 13,939 | AGD Type 2: 43,642 |
| Map resolution (Å; FSC=0.143) | Type 1: 3.4<br>Type 2: 3.4<br>Type 3: 8.3<br>PHF: 4.1<br>CTE Type 1: 4.7 | Type 2: 4.1<br>Type 3: 7.5<br>PHF: 3.7<br>SF: 3.4 | AGD Type 2: 3.6<br>AGD Type 3: 6.0<br>PHF: 9.4<br>SF: 8.8 | AGD Type 2: 4.2<br>AGD Type 3: 6.0<br>PHF: 4.3<br>SF: 4.4 | AGD Type 2: 3.2 | AGD Type 2: 3.4 |
| Helical rise (Å) | Type 1: 4.88<br>Type 2: 4.87<br>Type 3: 4.84<br>PHF: 2.42<br>CTE Type 1: 2.42 | Type 2: 4.85<br>Type 3: 4.84<br>PHF: 2.42<br>SF: 4.84 | AGD Type 2: 4.90<br>AGD Type 3: 4.84<br>PHF: 2.42<br>SF: 4.84 | AGD Type 2: 4.87<br>AGD Type 3: 4.84<br>PHF: 2.39<br>SF: 4.79 | AGD Type 2: 4.78 | AGD Type 2: 4.80 |
| Helical twist (°) | Type 1: -0.40<br>Type 2: -0.31<br>Type 3: -0.32<br>PHF: 179.4<br>CTE Type 1: 179.4 | Type 2: -0.36<br>Type 3: -0.4<br>PHF: 179.4<br>SF: -1.05 | AGD Type 2: -0.33<br>AGD Type 3: -0.38<br>PHF: 179.4<br>SF: -1.05 | AGD Type 2: -0.32<br>AGD Type 3: -0.38<br>PHF: 179.43<br>SF: -1.04 | AGD Type 2: -0.40 | AGD Type 2: -0.39 |

### Extended Data Table 4 | Cryo-EM data collection of FBD and FDD cases

| | FBD<br>Case1 | FDD<br>Case 1 |
|---|---|---|
| **Data collection and processing** | | |
| Electron Gun | XFEG | XFEG |
| Detector | Falcon III | Falcon III |
| Magnification | 75k | 75k |
| Voltage (kV) | 300 | 300 |
| Electron exposure (e–/Å$^2$) | 50 | 40 |
| Defocus range (μm) | 1.6-2.2 | 1.2-2.4 |
| Pixel size (Å) | 1.04 | 1.04 |
| Symmetry imposed | PHF: C1<br>SF: C1<br>CTE Type 1: C1<br>CTE Type 2: C1 | C1 |
| Initial particle images (no.) | 721,421 | 286,689 |
| Final particle images (no.) | PHF:<br>187,685<br>SF:137,235<br>CTE Type 1:<br>264,106<br>CTE Type 2: 24,182 | 214,400 |
| Map resolution (Å; FSC=0.143) | PHF:3.3<br>SF:3.2<br>CTE Type 1: 3.1<br>CTE Type 2: 3.7 | 3.2 |
| Helical rise (Å) | PHF:2.37<br>SF:4.74<br>CTE Type 1: 2.36<br>CTE Type 2: 2.36 | 2.36 |
| Helical twist (°) | PHF: 179.43<br>SF: -1.04<br>CTE Type 1: 179.42<br>CTE Type 2: 179.42 | 179.44 |
